# Recycler: an algorithm for detecting plasmids from *de novo* assembly graphs

**DOI:** 10.1101/029926

**Authors:** Roye Rozov, Aya Brown Kav, David Bogumil, Naama Shterzer, Eran Halperin, Itzhak Mizrahi, Ron Shamir

## Abstract

Plasmids are central contributors to microbial evolution and genome innovation. Recently, they have been found to have important roles in antibiotic resistance and in affecting production of metabolites used in industrial and agricultural applications. However, their characterization through deep sequencing remains challenging, in spite of rapid drops in cost and throughput increases for sequencing. Here, we attempt to ameliorate this situation by introducing a new plasmid-specific assembly algorithm, leveraging assembly graphs provided by a conventional *de novo* assembler and alignments of paired-end reads to assembled graph nodes. We introduce the first tool for this task, called Recycler, and demonstrate its merits in comparison with extant approaches. We show that Recycler greatly increases the number of true plasmids recovered while remaining highly accurate. On simulated plasmidomes, Recycler recovered 5-14% more true plasmids compared to the best extant method with overall precision of about 90%. We validated these results *in silico* on real data, as well as *in vitro* by PCR validation performed on a subset of Recycler’s predictions on different data types. All 12 of Recycler’s outputs on isolate samples matched known plasmids or phages, and had alignments having at least 97% identity over at least 99% of the reported reference sequence lengths. For the two E. Coli strains examined, most known plasmid sequences were recovered, while in both cases additional plasmids only known to be present in different hosts were found. Recycler also generated plasmids in high agreement with known annotation on real plasmidome data. Moreover, in PCR validations performed on 77 sequences, Recycler showed mean accuracy of 89% across all data types – isolate, microbiome, and plasmidome. Recycler is available at http://github.com/Shamir-Lab/Recycler

## Introduction

Plasmids are extra-chromosomal DNA segments carried by bacterial hosts. They are usually shorter than host chromosomes, circular, and encode nonessential genes. These genes are responsible for either plasmid-specific roles such as self-replication and transfer, or context-specific roles that can be beneficial or harmful to the host depending on its environment. Along with viruses and transposable elements, plasmids are members of the group termed mobile genetic elements [1] as they transmit genes and their selectable functions between microbial genomes. Plasmids play a central role in horizontal gene transfer [2], and thus genome innovation and plasticity - fundamental forces in microbial evolution.

Much interest has recently arisen for plasmid extraction and characterization, in particular because of their known roles in antibiotic resistance and in increasing metabolic outputs of agricultural or industrial byproducts. For instance, antibacterial resistance genes encoded on plasmids have long been known as a major issue for human health in clinical practice [3], but are also one of today’s standard tools in microbiology and genetics when used to select for specific cells [4]. In order to derive plasmid sequences (which may be known or novel), one may choose from the following approaches: sequence already isolated microbes with their residing plasmids, sequence the overall microbial community of genomes (termed metagenome) from some environment, or, as was recently described, sequence only the overall plasmid fraction from a given environment (termed plasmidome) [5],[6]. The first technique obtains a mixture of chromosomal and plasmid DNA occurring together in a single strain. Since sequenced reads are devoted to only a few different sequenced DNA elements (the genome in question or any of its mobile elements), each is expected to be highly covered, and thus for species having low repeat content a good assembly can be achieved.

For natural environments containing many elements, often including those that are difficult to culture [7] in a lab, metagenome assembly is attempted. This technique allows a much broader view of all taxa present and their plasmids, but is limited in that the characterization of each individual element depends on its coverage in the mixed DNA sample and the frequency of co-occurring repeats shared among different elements of the sample. Resulting assembled genomes of elements that are rare in the environment are thus often fragmented, and very high coverage [8] is needed for accurately assembling them. However, assembly of metagenomes remains a highly active area of research: current assembly outputs are lacking and do not represent the true genetic capacity and synteny of genomes present in complex microbial communities. Since most of the DNA in these environments is due to host genomes, this approach currently provides only limited resolution of plasmids.

Most recently, a third technique has emerged that allows recovery of far greater numbers of plasmids. Plasmidome sequencing [5], [6], [9] allows nearly all sequencing resources to be devoted to circular DNA. Using a protocol described in [5], chromosomal DNA is filtered out and circular DNA segments are selectively amplified. Based on this protocol, hundreds of new plasmids were identified in the cow rumen [6] and rat cecum [9]. In [9], Jørgensen et al. applied the protocol introduced in [5] combined with bioinformatic validation of circularity. This post-assembly analysis resulted in a 95% PCR validation rate out of 40 randomly selected assembled contigs. This success raises the prospect of *in silico* refinement of plasmids beyond the initial assembly. Although Jórgensen et al.’s method was shown to have a high validation rate, its output is limited by the contiguity of the underlying assembler’s contigs (in their case IDBA-UD [10]), because it provides no means of combining multiple overlapping contigs to form cycles. It is a filtering process meant to identify probable circular sequences among sequences already output by the assembler. To date, no tools for plasmid assembly from short reads have been introduced to address these limitations.

In all of the above approaches plasmid assembly is hindered by several inherent characteristics derived from their mobile nature. These characteristics include their tendency to carry repetitive elements such as insertion sequences and to share genes with other plasmids and microbial genomes. In the context of *de novo* assembly, repeats cause collapse of linear sequences sharing them as subsequences. This creates ambiguity in the sense that it becomes unclear which extensions entering the repeat should be paired with those exiting it, where sequences begin and end, and whether there are unique terminal points at all as opposed to the sequence being circular. *De novo* assembly for the sake of identifying plasmids can be augmented by long-read sequencing [11], [12] because such reads may be sufficiently long to bridge repeats short reads cannot. However, this approach is primarily limited to isolates or low complexity environments. This is evident in that long reads often depend on single molecule sequencing without amplification, thus only capturing relatively abundant DNA fragments. Besides repeats, chimeric sequences also present significant challenges to assembly, in that they create false connections between sequences and thus may lead to misassemblies.

Here, we wish to use more refined analysis in order to improve *de novo* assembly of sequenced plasmids. Our inputs are an assembly graph G = (V,E), and the mapping of paired-end reads responsible for the assembly to its nodes. The set of nodes V are sequences having associated lengths and coverage levels, and the set of arcs E is composed of directed connections among the nodes. Arcs are the result of branch points in the underlying de Bruijn graph: a branch node has outgoing arcs to two (or more) different nodes based on overlaps, and in many cases the assembler does not have a definite way of choosing which extension is true in order to simplify the branch into a linear path. We aim to generate a set of putative cycles that are likely to be plasmids, and assign a coverage level for each one.

After defining this problem formally below, we present an algorithm (and its implementation) designed to address it, called Recycler. Recycler leverages assembly graphs output by the SPAdes assembler [13] to specifically enable *de novo* assembly of plasmids. We show it greatly improves recovery of plasmids over naive assembly and alternative methods, namely Jórgensen’s and SPAdes’ built-in repeat resolution, introduced in [14]. We demonstrate Recycler’s performance by applying it on both simulated and real data. We find that Recycler greatly increases recall while maintaining high precision. This is established via comparisons performed on simulated plasmidomes of various sizes. We also show that Recycler can be applied for plasmid assembly on real data from a bovine rumen plasmidome and metagenome, and from two different Escherichia coli isolate strains. In the isolate cases, Recycler recovered most known plasmids, and predicted additional sequences that matched known mobile elements from different hosts – all of which were identical or nearly identical to known reference sequences. In all cases on real data, Recycler either matched or exceeded the proportion of outputs matching plasmid annotation, as described in [6].

### Related work

We note plasmid assembly is a multi-assembly problem, as described in the context of RNA-Seq transcriptome assembly [15]. Formulations of such problems often aim to generate a minimal set of paths that maximize agreement with observed data [15]–[17]. These methods usually employ network flow formulations, which admit polynomial-time algorithms for minimizing flow cost on the network; this flow corresponds to a convex function of the sum of coverage differences between observed and estimated coverage levels. However, these methods resort to heuristics in selecting a minimal set of paths to cover the entire graph, as splitting a flow into a minimal number of path and cycle components is an NP-hard problem [18].

Recycler does not aim to generate a set of paths explaining all coverage levels, and thus does not depend on a global objective function encompassing all nodes or edges. This approach is avoided because of the presence of linear paths due to either plasmids not fully covered during sequencing or bacterial host genomes housing plasmids, which may introduce noise into coverage levels observed and will not be part of the solution. Avoiding a global objective imposing parsimony on paths also allows Recycler to benefit from a polynomial time algorithm for generating ‘good’ cycles. Thus, Recycler’s approach is similar to StringTie [19], in that both repeatedly seek locally best paths or cycles and use coverage levels estimated on those to update coverage levels on the original graph, until some stopping criterion is met. We note the set of cycles desired is explicitly not minimal, as in cycle cover formulations [20]. For example, given a figure 8 component (Supplementary figure S1, I.) Recycler may represent it as two cycles separated by distinct coverage levels, where a minimal cover would use only one cycle. Instead, we wish to cover as much of the graph as possible with ‘good’ cycles.

## Methods

### Overview of Recycler

We present the input to our problem as a directed graph with vertices corresponding to non-branching sequence contigs and edges corresponding to connecting overlapping k-mers. (The graph can be viewed as obtained from the de Bruijn graph of order k of the sequence data by contracting edges (u,v) whenever u has outdegree 1 and v has indegree 1, and the sequence contig of the new node replacing u and v is the concatenation of their sequences). Each node has a coverage value reflecting its abundance in the input sequences. We search for cycles in the graph that will correspond to plasmids. Cycle sequence length, number of vertices, and coverage uniformity are factored in the selection process. We also use paired-end read mappings including mates on different nodes as a proxy for which of the nodes may have emerged from the same physical DNA fragment. This provides a means of inferring whether a candidate cycle is a plasmid or a genomic segment including repeats that lead to ambiguous cycles in the graph. Once a best cycle is selected, its latent coverage level is determined and subtracted from those of all participating nodes. Nodes whose resulting coverage values become non-positive are then removed from the graph, allowing only those with some remaining coverage the opportunity to take part in additional cycles. Hence, the whole process can be viewed as greedily “peeling off” cycles from the graph. Ideally, one would like the process to end in an empty graph, in which case the input graph would be exactly the union of the cycles found. In reality, the process is stopped when quality criteria for new cycles in the remaining graph are unmet.

### Notations and Definitions

Our input is a directed graph G = (V,E), where V is a set of linear sequences having either a branch-point or terminal k-mer at each end and no internal branch-points. E is the set of overlaps between nodes, where E = {(u,v): the (k-1)-mer suffix of u = the (k-1)-mer prefix of v }. We call a node in *G* simple if its indegree and outdegree are 1. A node v corresponding to sequence s of length l(s) is assigned two positive values, len(v) and cov(v). len(v) = l(s) – *k* + 1 is called the *length* of the node (the subtraction is in order avoid double-counting bases common to overlapping segments at their ends). cov(v), its *coverage*, reflects the average number of times each k-mer in s appears in the input read data. The input can be produced by a short read assembly tool. We further assign a *weight* 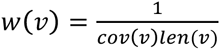 for each node v, resulting in low weight for high coverage and long nodes. Longer contigs tend to be less prone to random fluctuations in coverage, and are thus more reliable coverage indicators. For each cycle c in the graph, we assign each node *v* ∈ *c* a value *f*(*c, v*) representing its length fraction in 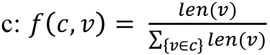. The value *f*(*c, v*) is used to define the mean and standard deviation of weighted coverage of cycle c as *μ*(*c*) = Σ_*v*∈*c*_ *f*(*c*,*v*)*cov*(*v*) and *STD*(*c*) = 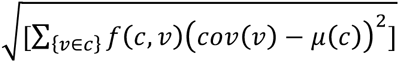 respectively, and consequently the coefficient of variation of c, *CV*(*c*) = *STD*(*c*)/*μ*(*c*) CV(c) is used to allow direct comparison of variation levels between cycles, independently of the magnitude of coverage of each. CV is indicative of coverage uniformity along the cycle, and plasmids are expected to have uniform coverage levels that in many cases are different from other plasmids and their hosts. Thus, a cycle with a lower CV is more likely to correspond to a plasmid than cycles with higher CV.

### Our approach

Intuitively, plasmids should form cycles that are distinctive from the rest of the graph and have near uniform coverage. We also expect plasmid cycles to include few nodes, as each additional node introduced for a fixed sequence length increases fragmentation and the tendency of nodes to be common to more than one path. With this in mind, we search for ‘good cycles’ in the graph that potentially correspond to plasmids. Formally, we define a good cycle as a simple cycle *c* in the graph satisfying the following constraints:

1. **Minimum path weight for some edge:** ∃(*u*,*v*) ∈ *c* such that *c* \ (*u*,*v*) (the path obtained by removing (*u, v*) from *c*) is a minimum weight path (by sum of weights w(v)) from v to u.
2. **Low coverage variation:** *CV*(*c*) ≤ *τ*/ |*c* |, where τ is a defined threshold and |*c* | is the number of nodes in *c*.
3. **Concordant read mapping:** For pair *r*_1_, *r*_2_ of paired-end mates, if *r*_1_ maps to a simple node in *c*, then *r*_2_ must also map to some node in *c*.
4. **Sufficient sequence length:** Σ_*v* ∈ *c*_ *len*(*v*) ≥ *L*, where *L* is a defined threshold.

The first constraint is critical in order to avoid merging of two or more plasmids that are connected through a repeated region (supplementary figure S1, panel I.). In addition, lower weight cycles correspond to longer sequence length and higher coverage nodes, and tend to have fewer nodes. Furthermore, for each edge this constraint uniquely determines at most one cycle that passes through the edge, thus avoiding consideration or enumeration of an exponential number of possible cycles. We note there are special cases allowing for cycles that visit a single node more than once; such a case is shown in supplementary figure S1, panel II. The second constraint ensures that the coverage variation is low, thus again increasing our confidence that the cycle corresponds to exactly one plasmid. Moreover, this constraint implicitly ensures high coverage cycles, since low coverage cycles tend to have higher CV value. The third constraint exploits paired-end reads. Suppose we have a read pair *r*_1_, *r*_2_ and *r*_1_ maps to a certain node in the candidate cycle *c*. We expect *r*_2_ to map to the same cycle, unless *r*_1_ falls on a node that is common to *c* and some other path *p* overlapping with it. In that case *r*_2_ may map to *p* as well. Simple nodes are less likely to overlap with several cycles and path, and the third constraint leverages this observation. We waive this constraint in case the coverage of *c* is sufficiently high, as in such cases the cycle “stands out” from the background coverage. See Supplement for details.

The above definition of a good cycle provides a mechanism for the identification of putative plasmids. Recycler processes each strongly connected component separately. It repeatedly finds a good cycle with minimum CV value, assigns it *latent coverage* equal to the mean cycle coverage and subtracts that coverage from the graph, creating a new *residual coverage* (Figure 1). The weights of the vertices in the cycle are updated based on their new coverage values, and vertices whose resulting coverage values become non-positive are removed from the graph, allowing only those with positive residual coverage the opportunity to take part in additional cycles. After each such change, cycles are recalculated the same way using the updated coverage levels. This process continues as long as new good cycles are found. To avoid examining a potentially exponential number of cycles, we consider one minimum weight cycle through each edge in the graph. Algorithm 1 sketches the procedure for a single component. See the supplement for additional details.

**Figure 1.**
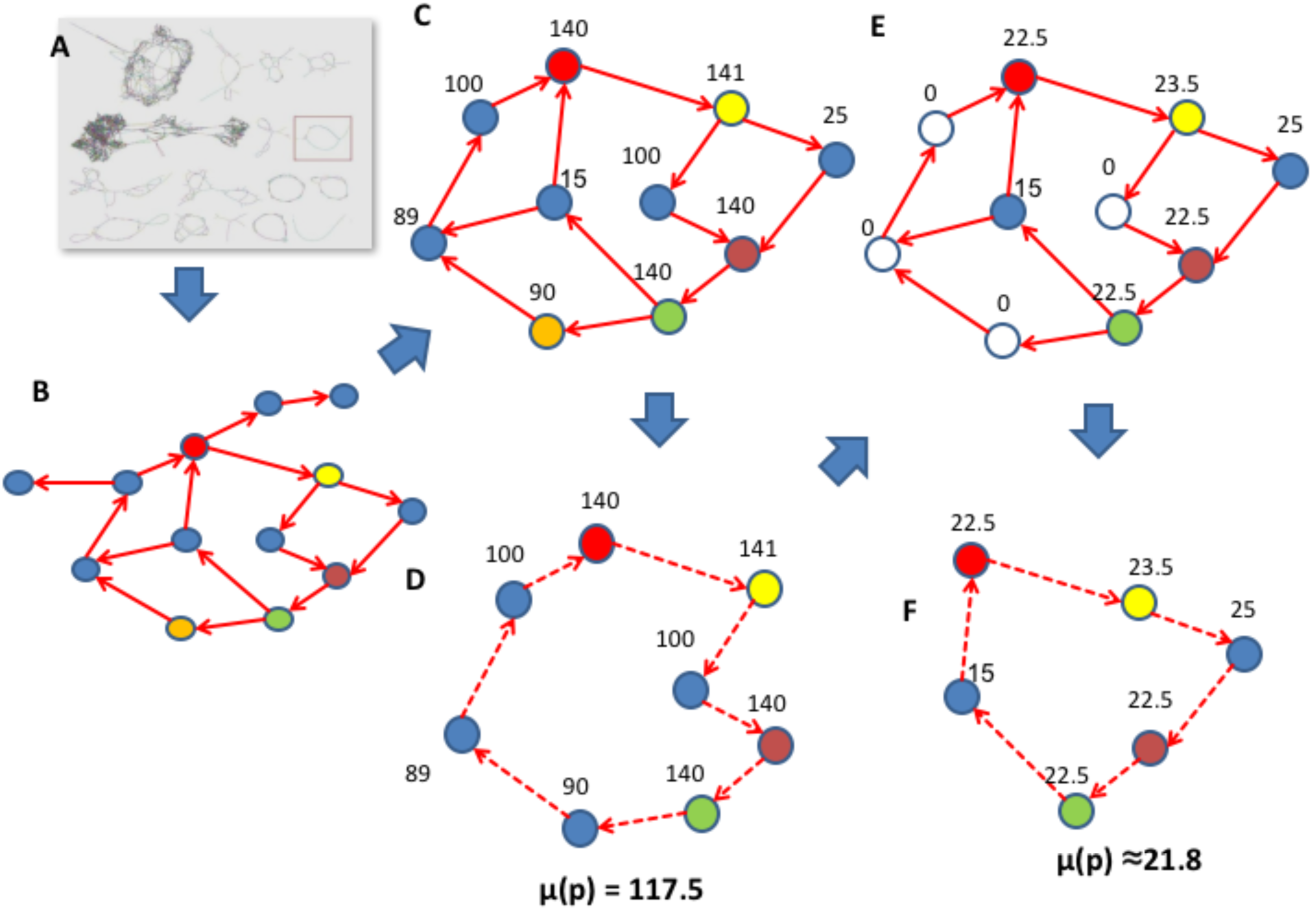
Recycler work-flow. **A.** The assembly graph. **B.** A single component is selected from the assembly graph (framed in A) and represented with vertices for contigs (arbitrarily colored for identification) and edges for connecting k-mers. **C.** The reduced component after tip removal. Vertex values are observed contig coverage. For simplicity, all lengths are assumed to be equal and not shown. **D.-F.** Two successive steps of peeling cycles are shown with their respective latent coverage assignments. Uncolored vertices correspond to contigs with zero coverage that are removed.

#### Algorithm 1

( Inputs: *G* = (*V, E, len, cov, W*), *τ, L*; Output: Σ, the set of cycles )

*Compute shortest cycles passing through each edge*,

For each edge (u,v)

Compute a minimum weight path p from v to u, if one exists

Compute the CV of the cycle (p,(u,v))

Return the set of cycles S

While Σ changes,

Compute a set S of shortest cycles passing through each edge

Consider each cycle c in S in increasing order of CV values

If c is good and not in Σ

Add c to the solution set Σ

Compute the latent coverage level of c

Update the residual coverage of all cycle nodes, removing

nodes with non-positive residual coverage

Output Σ and end

### Complexity

Algorithm 1 presented above terminates in polynomial time. In each iteration, if any good cycles exist, one is chosen and its mean coverage is calculated. There is at least one node in the cycle with coverage smaller than the mean coverage of the cycle, which is subsequently removed from the graph. Therefore, in each iteration at least one node is removed, and the number of iterations is bounded by the number of nodes. Using Johnson’s algorithm [21], the runtime of each iteration is *O*( |*V* |^2^log |*V* | + |*V* | |*E* |). Running times are further reduced by computing the strongly connected components of *G* and working separately on each one.

### Generating simulated plasmidomes

We simulated error-free paired-end reads from plasmids using BEAR [23], a read simulator designed to generate artificial metagenome data. To avoid introducing coverage drops at sequence ends typical of linear sequences, we modified BEAR [https://github.com/rozovr/BEAR] to allow sampling of reads bridging reference sequence ends, as is observed for circular sequences. Plasmid reference sequences were selected from the NCBI plasmids database and from plasmid sequences reported in [6], filtered to include 2760 sequences with a length range of 1 to 20 kbp with a mean of 6337 bp. Five datasets were created, composed of 100 bp mates (read pair ends), with insert sizes ~ *N*(500,100), varying from 1.25M pairs sampled on 100 reference sequences doubling successively up to 20M pairs sampled on 1600 sequences. Abundance levels were assigned using BEAR’s low complexity option, which concentrates high abundance to few species using a power function with parameters derived from [24]: the function takes the form ci^d^, where c=31.4 and d=-1.28, and i is iteratively assigned values from 1 to the number of species simulated. These values are then normalized by their sum to yield a probability distribution.

### Evaluating performance

To test recovery of the ground truth sequences by each plasmid detection program, we used the Nucmer alignment tool [25], which is designed for efficiently comparing long nucleotide sequences such as those of whole plasmids or chromosomes. In order to simplify this process, we modified reference sequences to remove non-ACGT characters before read simulation and alignments. To avoid fragmented alignments caused by differences in start positions, we concatenated each reference sequence to itself before mapping; this allowed identification of complete matches at the center of the concatenated contigs when they were present. Output cycles of each tested program were defined as true positives (TP) if they had 100% identity hits covering at least 80% of one of the reference sequences. False positives (FP) were any output cycles not meeting these criteria, and false negatives (FN) were reference sequences not aligned to in the output set using these criteria. Based on these conventions, 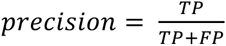 and 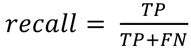. We used the score 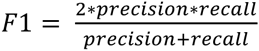 [26] to combine these measures in a manner that weighs precision and recall equally.

### Primer design and PCR validation of plasmid contigs

The plasmidome dataset was divided into two separate subsets, including simple (single node) cycles (N=370) and complex (multi-node) paths within the graph (N=50). Each of these was divided into coverage bins, and selected representatives from each bin (High coverage –60-1000x, mid-high coverage – 15-60x, mid-low coverage – 5-15x, low coverage – 1-5x) were validated by PCR. Overall, 24 simple cycles and 39 complex cycles were chosen for PCR validation. From the metagenome dataset (N=40), all assembled plasmids were of the same coverage bin (1-5X) from which 10 plasmids were randomly selected for validation. This was also the case for the E. coli E2022 isolate (N=4) for which all plasmids were validated by PCR, aside from a recovered Phi X control sequence. Primers were designed to produce an amplification product only if their template is circular; this was achieved by directing the opposing primers towards the edge of the linear plasmid contig. PCR reactions were carried out using Advantage GC Genomic LA PCR Polymerase (Clontech) according to the manufacturer’s instructions. The PCR reactions were as follows: 1.5 μl Advantage buffer (X10), 0.6 μl of each primer (5mM), 0.15 μl Ex Advantage GC Genomic LA DNA Polymerase, 100 ng of template DNA, 1.5 μl of dNTPs (10mM) and DDW was added to a final volume of 25 μl. All PCR reactions were carried out in a Sensoquest thermocycler (Gottingen, Germany).

## Results

We first simulated plasmidomes using known references. We used these data sets to assess Recycler’s precision and recall (along with those of alternative methods) by comparing predictions against the ground truth known by the simulation design. We also tested Recycler on real data from two *E. Coli* isolates, and both a cow rumen metagenome and plasmidome[6]. For the bacterial isolates that have been sequenced, predicted plasmids were compared against the reference sequences directly. Since no references are available for metagenome and plasmidome data, we evaluated the accuracy by PCR validation [9] and by measuring the proportion of predicted plasmids having proper annotation as done in [6]. Recycler’s inputs were assembly graphs generated by SPAdes version 3.6.2 [13], and alignments generated by BWA version 0.7.5 [22].

### Simulated plasmidomes

We simulated paired-end reads from known plasmids, and created five datasets of 100, 200, 400, 8000 and 1600 plasmids. Plasmid abundance was distributed so that few plasmids have high abundance. Dataset sizes were 1.25, 2.5, 5, 10 and 20M pairs, respectively (see Methods for details). Each such dataset was assembled with SPAdes and subsequently its output contigs and assembly graphs were used as inputs to the tested methods. Recycler was compared with SPAdes with and without repeat resolution (RR), and to a simplified version of Jórgensen’s method (described in the appendix). We used SPAdes’ outputs before the repeat resolution stage as inputs to Recycler and to a Jórgensen’s method, as we found that contigs have greater precision before RR when compared to reference sequences (as shown in supplementary Table 1). The mapping results are presented in supplementary Table 1 and Figure 2.

**Figure 2.**
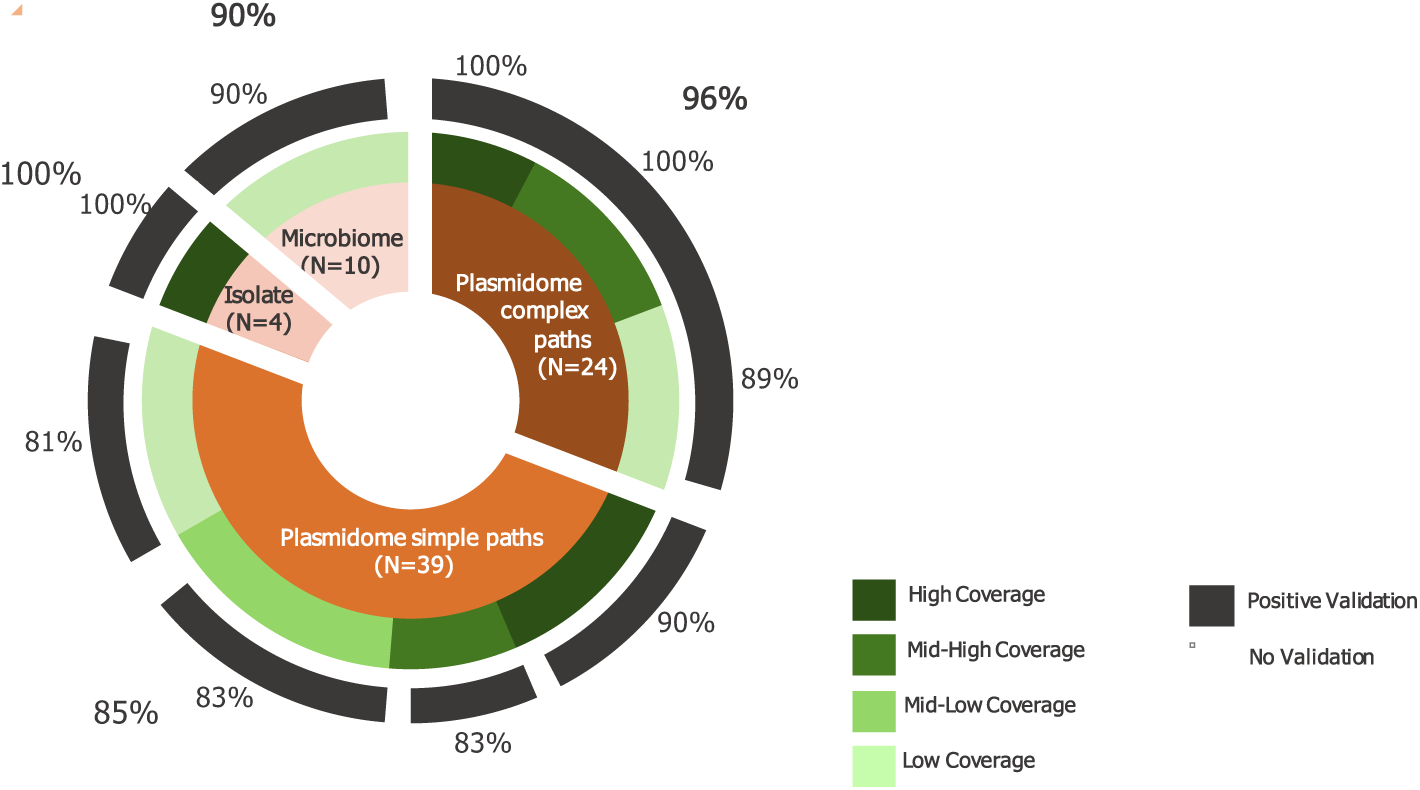
PCR based validation of Recycler’s plasmid predictions. The inner most ring (brown) describes the different datasets for which PCR validation was carried out. The second ring (green) describes the distribution of tested coverage levels among plasmids tested. High coverage: 60-1000x, mid-high: 15-60x, mid-low: 5-15x, low: 1-5x. The black lines indicate the success rate (written above each bar) in PCR validation in each category. Finally, the numbers outside the light orange lines indicate the overall success rate for each dataset. Positive validations are those confirming the presence of predicted circular sequences (see text).

**Figure 3.**
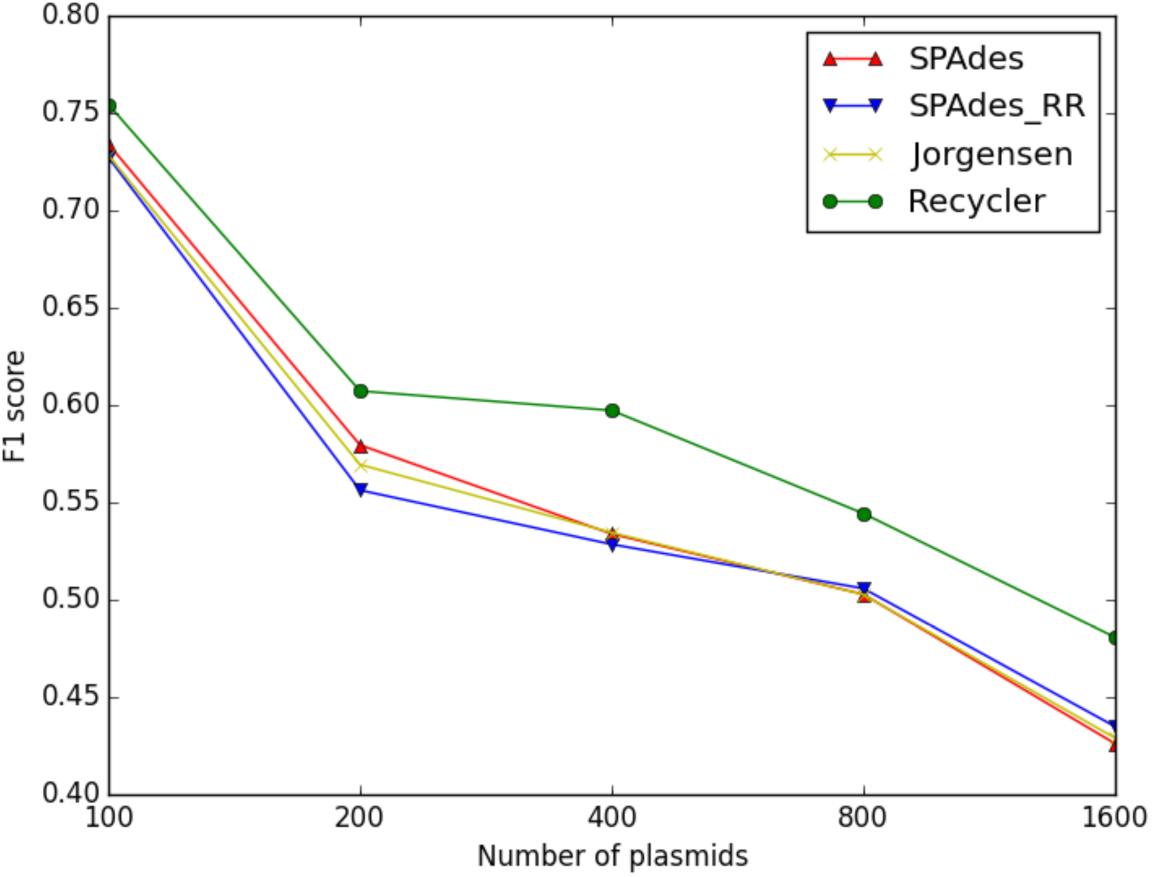
Methods performance on simulated data. Results are shown for SPAdes without repeat resolution (RR), SPAdes with repeat resolution, the method of Jórgensen et al, and Recycler. The contigs of SPAdes before RR were used as input for the three other methods. Recycler also relied on the graph produced at this stage. F1 score calculation is described in the main text. The x axis shows the number of simulated reference sequences in each case.

**Figure 4.**
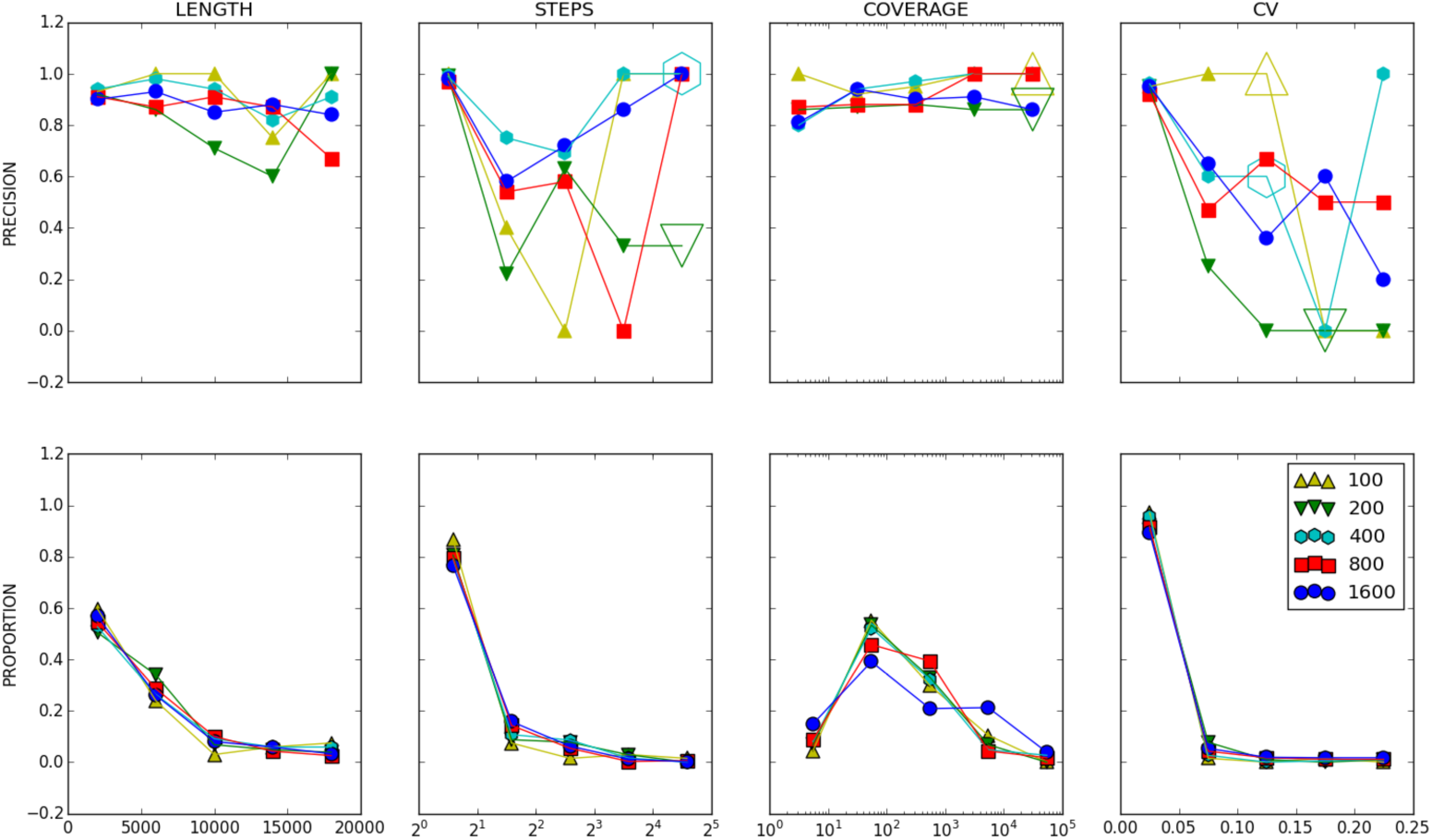
Recycler’s precision, stratified by different properties. TOP: For simulated reads, true positive (100% identity over 80% reference length) alignment proportions were tallied inside 5 bins corresponding to value ranges of different properties - total assembly length (LENGTH), number of nodes in the cycle (STEPS), mean coverage level on the paths (COVERAGE), and cycle coefficient of variation (CV). Each point represents the precision rate for all simulated plasmids included in that range in the specified reference set. Reference sets are denoted by different colors and marker shapes. Enlarged empty markers are used to indicate the absence of any instances having the given property & bin combination. BOTTOM: relative proportions of counts inside each bin out of all outputs.

**Table 1.**
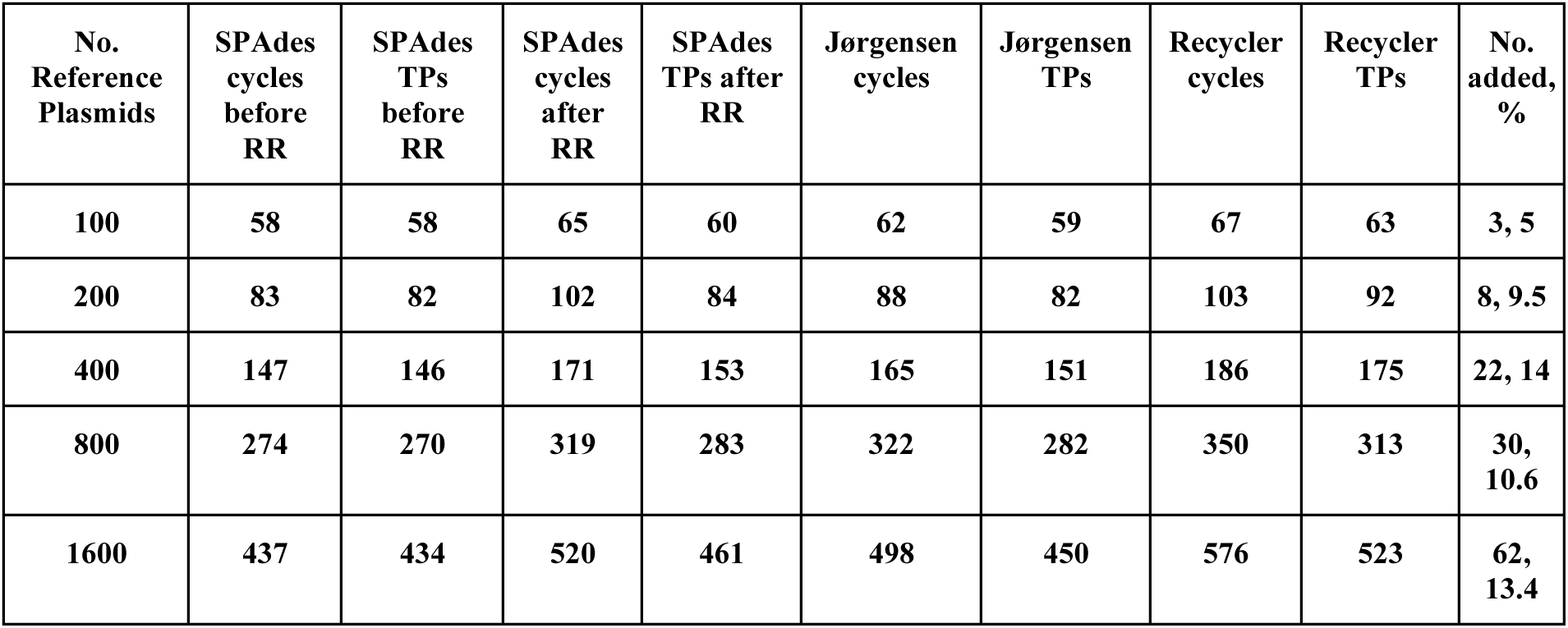
Counts of output cycles vs. true positives (TPs) for each method. The number of reference plasmids reflects the number of sequences that were simulated. The number of cycles reflects the total output by each method, and the true positives represent those that match some reference sequence based on alignment criteria defined in the main text. The last column shows Recycler’s advantage over SPAdes with repeat resolution in number of plasmids and percentage added.

**Table 2.** – Real data results summary

|  | E2022<br>Recycler | E2022<br>SPAdes | JJ1886<br>Recycler | JJ1886<br>SPAdes | metagenome<br>Recycler | metagenome<br>SPAdes | plasmidome<br>Recycler | plasmidome<br>SPAdes |
| --- | --- | --- | --- | --- | --- | --- | --- | --- |
| No. seqs annotated as plasmids | 5 | 3 | 7 | 3 | 12 | 10 | 249 | 203 |
| No. seqs with hits on [9] data | 0 | 0 | 0 | 0 | 1 | 0 | 53 | 50 |
| No. seqs with any nr annotation | 5 | 3 | 7 | 3 | 37 | 30 | 314 | 264 |
| total No. cys | 5 | 3 | 7 | 3 | 40 | 33 | 420 | 361 |
| % annotated cys as plasmids | 100 | 100 | 100 | 100 | 32 | 33 | 79 | 77 |
| % annotated cys as plasmids or [9] | 100 | 100 | 100 | 100 | 35 | 33 | 96 | 96 |
| % annotated at all | 100 | 100 | 100 | 100 | 93 | 91 | 75 | 73 |

**Table 3.** – plasmids detected by Recycler on E. Coli strain JJ1886

| Reference seq. length (Kb) | Output seq. length (Kb) | Top blast hit | ID (%) | Len. (%) |
| --- | --- | --- | --- | --- |
| 1.6 | 1.6 | JJ1886 plasmid | 100 | 100 |
| -- | 1.6 | *S. Aureus* plasmid | 100 | 100 |
| -- | 2.2 | *S. Aureus* plasmid | 100 | 100 |
| -- | 2.4 | *S. Chromogenes* plasmid | 100 | 100 |
| 5.2 | 5.2 | JJ1886 plasmid | 100 | 100 |
| 5.6 | 5.6 | JJ1886 plasmid | 100 | 100 |
| 56.0 | 56.0 | JJ1886 plasmid | 100 | 100 |
| 103 | -- | -- | -- | -- |

**Table 4.** – plasmids detected by Recycler on *E. Coli* strain E2022

| Reference seq. length (Kb) | Output seq. length (Kb) | Top blast hit | ID (%) | Len. (%) |
| --- | --- | --- | --- | --- |
| -- | 1.5 | *E. Coli* FHI63 plasmid | 100 | 100 |
| 2.2 | 2.1 | *E. coli* GXEC6 plasmid | 100 | 100 |
| 4.1 | 4.1 | Jørgensen et al. plasmid | 100 | 99 |
| -- | 5.4 | Enterobacteria phage phi-x174 | 100 | 100 |
| 35 | 33.1 | *E. Coli* GXEC6 plasmid | 97 | 99 |
| 98.3 | -- | -- | -- | -- |
| 103 | -- | -- | -- | -- |

As expected, recall generally decreased as the number of simulated plasmids increased. This was common to all tested methods. In general, we found that Recycler generated more predictions than other methods, leading it to have higher recall than alternative approaches while maintaining high (~90%) precision. The net performance effect is shown in Figure 2 and Table 1 in the supplement: Recycler maintains the lead in all cases with 5-14% advantage in both F1 and fraction of true positives. We also found that the number of additional Recycler true positives over those provided by SPAdes generally increased with higher complexity; this culminated in Recycler adding 62 (13%) true positives to SPAdes’ output on the 1600 plasmid set (523 vs. 461).

To further characterize Recycler’s performance, we categorized its predictions in terms of mean total cycle length, number of segments in the cycle (“steps”), cycle coverage, and CV value calculated at the stage the cycle was removed. For each category, values were subdivided into five ranges. In Figure 3, we show the precision values and the relative proportions of counts in the specified ranges. Based on this stratification, it can be seen in Figure 3 that Recycler shows little dependence on mean coverage or length, but does often preclude candidate cycles that have high CV values or number of steps. This is reflected in the sharp drop-off in the plots as the number of steps or the CV grows.

## Real data

All of Recycler’s results on real data were subjected to quantification of annotation results as described in [6] and compared against cycles present in the output produced by SPAdes. These results are detailed below and a summary of them can be found in Table S2 in the Appendix.

### Circular integrity of assembled plasmids

Overall, 89% of the 77 chosen plasmids were validated as circular DNA molecules. The predicted plasmids from the different samples did not differ in the success rate of circular validation. As coverage has a key role in de novo assembly and Recycler’s performance, we wished to measure whether the integrity of assembled plasmids would be affected by varying mean k-mer coverage. To this end, we validated circularity of plasmids of different coverage levels ranging from 1x -1000x divided into bins. As can be seen in Figure 1, there was a slightly lower success rate for the lower coverage plasmids. However, coverage and validation rate were not found to be significantly correlated. Additionally, the high number of predicted plasmids in the plasmidome data set allowed us to measure the effect of the “complexity" of the path on the graph on the integrity of the plasmids. When more edges are involved in a cycle, it is more complex, and the chance of noise in coverage levels and errors in sequence increases. Thus, we divided this dataset into two bins according to path length on the graph: simple: single node (self-edge) paths, complex: two nodes or more. These two bins did not show difference in their validation rate, further stressing Recycler’s strength in extracting plasmids from complex paths.

### *E. Coli* isolate data

We ran Recycler on two *E. Coli* strains: JJ1886, downloaded from http://www.ebi.ac.uk/ena/data/view/SRX321704. and E2022, sequenced locally. Annotation for plasmids found in both strains was provided in [27]; comparisons against Recycler outputs with this annotation are reported in Supplementary Tables 3 and 4. Of the five plamids known for JJ1886, Recycler output four complete matches (100% identity over 100% length) having lengths 55.9, 5.6, 5.2, and 1.6 kbp. It also output three additional sequences which completely matched previously reported plasmids: two are known to be present in *S. aureus*, and one in *S. chromogenes*. Further tests will be needed in order to validate whether these additional hits are truly present in the sequenced sample, and furthermore, whether they are stable residents of the tested hosts or were present as a result of contamination. When tested on E2022, Recycler performed similarly, recalling most of its known plasmids and outputting a few additional cycles that were complete or near complete matches to known plasmids and one phage. These results are also presented in Supplementary Table 2. In summary, all reported isolate hits represent highly accurate matches to known mobile elements, and most known plasmids for these strains were recovered. In both cases Recycler missed the longest known reference plasmids; it remains to be seen whether this is due to Recycler’s use of a shortest path formulation, lack of significant coverage difference between these plasmids and the host genome, or other factors.

### Plasmidomes data

A bovine rumen plasmidome sample was prepared as described in [6]. This data consisted of 5.1 M paired-end 101 bp reads (trimmed to varied sizes for the sake of adapter removal) with an expected insert size of 500 bp [data available upon request]. Recycler output 420 cycles when provided this data. According to ORF prediction performed as in [6], 314 of the 420 had significant annotation hits. 96% of those matching annotations either matched plasmid annotations or aligned with plasmids reported in [9]. Thus, a majority are likely to be plasmids.

### Metagenome data

Metagenome data was derived from the rumen of a different cow residing in te same stable as the cow used to derive the plasmidome data. This data consisted of 7.5 M paired end 150 bp reads with expected insert size of 500 bp [data available upon request]. Recycler produced 40 cycles when run on this data. According to ORF prediction, 37 of the 40 had significant annotation hits. 35% of those matching annotations either matched plasmid annotations or aligned with plasmids reported in [9]. The proportion of reported cycles matching known plasmid annotations was slightly higher than for simple cycles output by SPAdes (33%). Overall, this test reflects the trend seen elsewhere [8] of weak annotation results emerging from metagenome assembly of highly diverse environmental samples.

## Discussion

In this article, we describe Recycler, a new algorithm and the first tool available for identification of plasmids from short read-length deep sequencing data. We demonstrate that Recycler discovers plasmids that remain fragmented after *de novo* assembly. We have adapted the approach of choosing among likely enumerated paths using coverage and length properties, (often applied in transcriptome assembly, e.g., [16], [19], [17]) for extracting a specific but common inhabitant of metagenomes. We showed that many more real plasmids can be found by only generating likely cycles on the assembly graph versus alternative methods. We validated this approach on both real and simulated data.

Recycler displays high recall and precision on simulated plasmidomes, and we have developed a means of separating real plasmids from cycles due to repeats in isolate data. As we have noted, coverage can be very useful for the latter, but the assumption that coverage will always differ significantly between plasmids and their host genome does not hold universally. It is worth noting that as new plasmids are identified and their common sequence motifs are observed, both reference-based identification and a priori trained prediction of plasmid features can be improved and harnessed for supplementing identification based on coverage and length features alone. We aim to investigate how such knowledge can be leveraged for increased precision without sacrificing recall.

Further investigation will be needed to assess how plasmids can be extracted from environmental samples, in spite of the limitations now hampering metagenome assembly. This is currently challenging, as diverse genomes require very high coverage for rare species to be captured, but such high coverage data demand computational resources beyond reach of most investigators. While new techniques have aimed to address this problem [8], [28], they have yet to see widespread use, and work best when paired with multiple samples to allow for species separation by co-abundance signatures. Along with addressing these concerns, it remains to be seen whether a mixed approach of pre-screening environmental samples for plasmids and computationally filtering them out may benefit metagenome graph simplification.

## Acknowledgements

Research partially supported by the Israel Science Foundation grants no. 1425/13 (EH), 317/13 (RS), 1313/13 (IM) and the ISF-NSFC joint program 2015-18 (RS). Additional support was provided by the European Research Council (ERC) under the European Union’s Horizon 2020 research and innovation program (grant agreement No 640384, IM). RR was supported in part by a fellowship from the Edmond J. Safra Center for Bioinformatics at Tel Aviv University, an IBM PhD fellowship, and by the Center for Absorption in Science, the Israel Ministry of Immigrant Absorption. EH is a Faculty Fellow of the Edmond J. Safra Center for Bioinformatics at Tel Aviv University. RR wishes to thank Kobi Perl and David Pellow for helpful comments given in the preparation of the manuscript.

## Appendix

**Figure S1.**
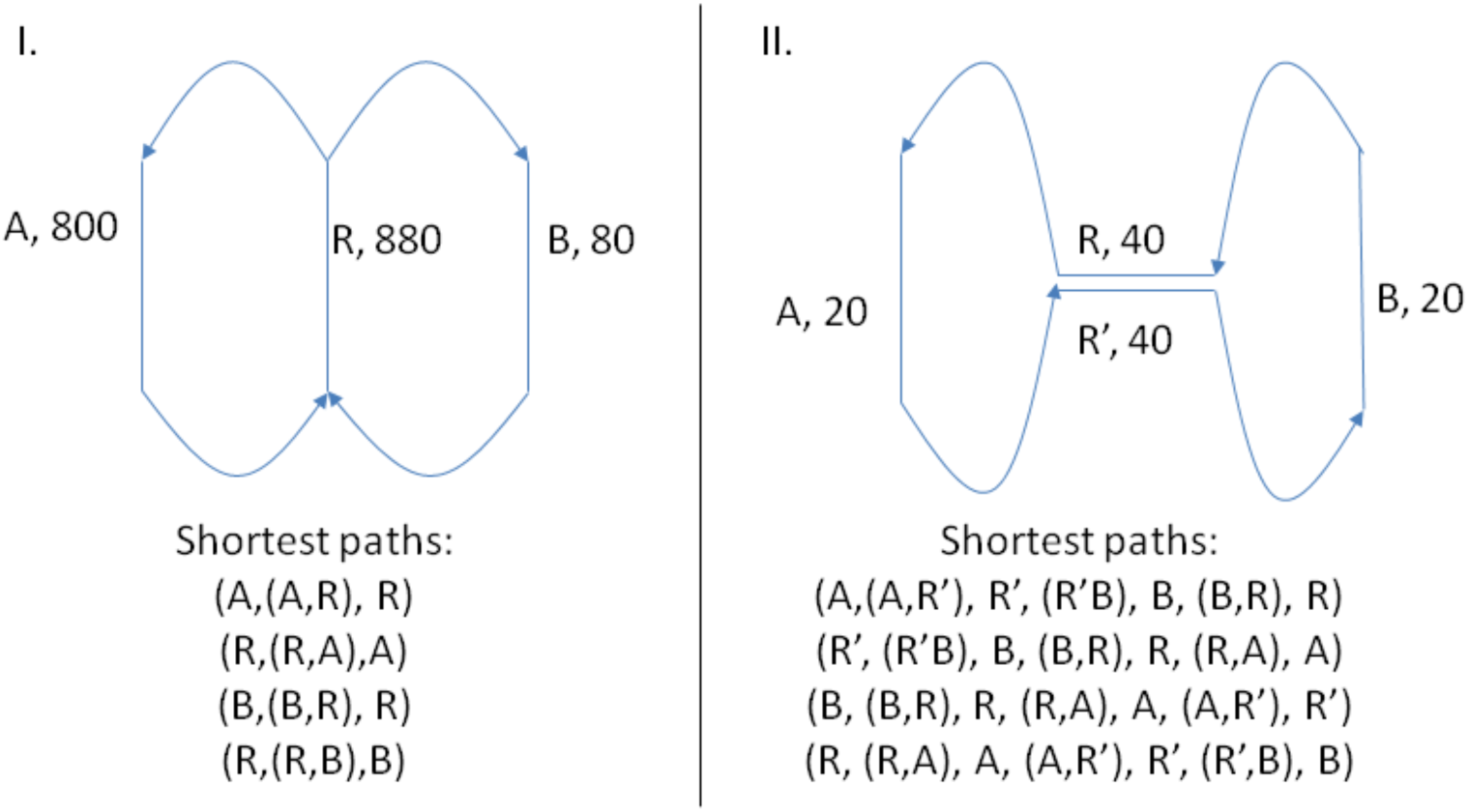
Shortest paths through repeats. Nodes are represented as lines, and edges as arrows. For each node x, x’ represents its reverse complement node. All node lengths are 1. I.) Recycler will consider only the shortest paths shown as candidates. The shortest path criterion described in the Methods section allows Recycler to avoid looping in a path with the shape “figure 8.” II.) For the component on the right, Recycler must consider shortest paths including the figure 8 path covering the entire component, as R and R’ have different adjacencies. In such cases, Recycler will assign equal parts of the coverage to R and R’: each gets 20. This demonstrates the use of the shortest path criteria in only allowing repeated traversal of nodes when warranted by coverage and orientation.

### Jørgensen’s method

We only used the first part of the protocol described Jørgensen’s method in order to allow for maximal recall; the second part involved further filtering (and thus reduction) of the first part’s results. Circular contigs were identified by finding those having opposite ends that overlap. These were then refined by breaking those that do have such overlaps into halves, and then gluing the far ends by applying the minimus2 assembler, part of the AMOS package.

**Tables 3, 4: Recycler output sequences compared to known reference sequences of two E. Coli strains. Values shown are sequence lengths and percentages of sequence identity (ID) and reference sequence sequence length covered (Len). Cells marked with ‘—‘ reflect unmatched sequences - either Recycler output a sequence not present in the reference set, or a sequence in the reference set wasn’t found by Recycler.**

### Algorithm details

**Definitions, notation** For all 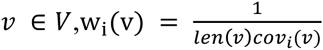. These are used to find minimal cost paths in calls to Johnson’s [21] algorithm. When applying Johnson’s algorithm, each edge (u,v) is assigned weight w(u), as Johnson’s algorithm depends on edge weights instead of node weights to calculate minimum cost paths. Each time a cycle c is peeled from a component, coverage levels are updated to be *cov*_*i*+1_ = max (*cov_i_* – *μ*(*c*), 0) for all nodes in c.

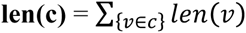

|**c** | = the number of contigs in c

**thresh** is set to the 75^th^ percentile of simple node coverage levels for plasmidome data sets, and the 95^th^ percentile for isolate data sets. This threshold is used in the testing for good paths. We use a different threshold for the two scenarios as it is easier to distinguish between coverage levels typical for the genome vs. foreign elements on isolate data than it is for more diverse environmental samples.

**‘Good’ path conditions:**

– len(c) ≥ *ℓ*_{*min* }_
– 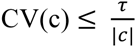
– CASE_A: All reads *r*_*i*1_ mapping to simple nodes in path c have their mates *r*_*i*2_ mapping to some *v_i_* such that *v_i_* is on c
– CASE_B: μ(c) ≥ thresh

Define (***r***_*i*_, ***v***_*i*_) as an alignment of read r_i_ to v_i_. Then the set of paired-end reads with mates aligned to different contigs is denoted as **A_*GR*_** = {(***r***_**1**_, ***v***_**1**_), (***r***_**2**_, ***v***_**2**_): ***r***_**1**_ = ***mate***(***r***_**2**_), ***v***_**1**_, ***v***_**2**_ ∈ ***V***, ***v***_**1**_! = ***v***_**2**_ } Define SCC(G) to be the set of strongly connected components of G

#### Algorithm

<u>Inputs</u>: *G* = (*V,E*), *τ, ℓ_min_*, A_GR_

func **recycle**(G, A_GR_, *τ, ℓ_min_*),

cycles = {(u,u) : u ∈ *V* is a simple node }

for COMP in SCC(G),

cycles = cycles U **peel_cycles**(COMP, A_GR_, *τ, ℓ_min_*)

return cycles

func **peel_cycles**(COMP, A_GR_),

paths = **get_shortest_pred_paths**(COMP)

last_path_count = 0

last_node_count = 0

while (|final_paths | ≠ last_path_count OR |V_COMP_ | ≠ last_node_count),

last_node_count = | V_COMP_ |

last_path_count = |final_paths |

paths = sort_by_CV(paths)

curr_path = paths.pop() # get lowest CV path

if **is_good_path**(curr_path, A_GR_,) and curr_path ∉ final_paths,

**update_path_coverage_vals**(curr_path, COMP)

final_paths = final_paths U {curr_path }

paths = **get_shortest_pred_paths**(COMP)

return final_paths

func **get_shortest_pred_paths**(comp),

paths = { }

shortest_paths = Johnson(G,W)

for v in V_comp,

preds = {u: (u,v) ∈ E_comp }

(path, path_cost) = shortest_paths(v,u)

if path_cost < ∞,

paths = paths U {path }

return paths

func **is_good_path**(path, A_GR_),

if (len(path) ≥ *ℓ_min_*) AND ( (IS_CASE_A(path)) OR (IS_CASE_B(PATH, A_GR_))),

return TRUE

else,

return FALSE

func **update_path_coverage_vals**(path, G),

μ = weighted_mean_coverage(path)

for v in path,

cov(v,G) = cov(v,G) – μ

if cov(v,G) ≤ 0,

remove(v,G)

